# Is dopamine D1 receptor availability related to social behavior? A positron emission tomography replication study

**DOI:** 10.1101/218420

**Authors:** Pontus Plavén-Sigray, Granville J. Matheson, Petter Gustavsson, Per Stenkrona, Christer Halldin, Lars Farde, Simon Cervenka

## Abstract

**Background:** Associations between dopamine receptor levels and pro- and antisocial behavior have previously been demonstrated in human subjects using positron emission tomography (PET) and self-rated measures of personality traits. So far, only one study has focused on the D1-dopamine receptor (D1-R), finding a positive correlation with the trait social desirability, which is characterized by low dominant and high affiliative behavior, while physical aggression showed a negative correlation. The aim of the present study was to replicate these previous findings using a new independent sample of subjects.

**Methods:** Twenty-six healthy males were examined with the radioligand [^11^C]SCH-23390, and completed the Swedish universities Scales of Personality (SSP) which includes measures of social desirability and physical trait aggression. The simplified reference tissue model with cerebellum as reference region was used to calculate BP_ND_ values in the whole striatum and limbic striatum. The two regions were selected since they showed strong association between D1-R availability and personality scores in the previous study. Pearson’s correlation coefficients and replication Bayes factors were then employed to assess the replicability and robustness of previous results.

**Results:** There were no significant correlations (all p values > 0.3) between regional BP_ND_ values and personality scale scores. Replication Bayes factors showed strong to moderate evidence in favor no relationship between D1-receptor availability and social desirability (striatum BF01 = 12.4; limbic striatum BF01 = 7.2) or physical aggression scale scores (limbic striatum BF01 = 3.3), compared to the original correlations.

**Discussion:** We could not replicate the previous findings of associations between D1-R availability and either pro- or antisocial behavior as measured using the SSP. Rather, there was evidence in favor of failed replications of associations between BP_ND_ and scale scores. Potential reasons for these results are restrictive variance in both PET and personality outcomes due to high sample homogeneity, or that the previous findings were false positives.

## Introduction

The dopamine system is involved in a wide range of behavior. A series of molecular imaging studies suggest that regional levels of the D2-dopamine receptor (D2-R) in the human brain are negatively related to pro-social behavior, such as the personality trait social desirability (1–4), although a null finding has also been reported (5). Social desirability reflects how a person represents herself in a social setting in order to gain approval by others, and combines low dominance and high affiliation traits (6). Compared to the D2-R, research on the relationship between personality traits and the D1-receptor (D1-R), which show different intracellular mechanisms and brain distribution (7, 8), has been much more scarce.

In a previous publication (9) we reported a positive correlation between D1-R levels in striatum and social desirability in healthy subjects, while the opposite pattern was shown for measurements of aggressive personality traits. This finding mirrors that from animal literature (10) and when taken together with previous PET studies (1–4), suggests opposite regulatory mechanisms for the D1 and D2 dopamine systems in mediating pro- and antisocial behavior in humans. This in turn could have wide implications for diagnosing and treating psychiatric conditions associated with dysfunctional social behavior, such as antisocial personality disorder or social anxiety. However, common problems with studies using positron emission tomography (PET) to examine personality traits are that they often are based small samples (with risks of selection bias and non-normal or restricted variability), employ many outcome measures, and allow for flexible modelling options, which can increase the risk for false positive findings. It is therefore important to replicate findings using independent samples in order to assess the robustness of published results.

The objective of the present study was to perform a replication of our previously reported associations (9) between D1-R in the striatum and social desirability and physical aggression, using a new and independent sample.

## Materials and Methods

### Subjects and personality measures

Twenty-six male subjects (mean age=26.2 *±* 3.2) were recruited and participated in PET examinations with the D1-R radioligand [^11^C]SCH-23390 (11). Exclusion criteria were historical or present episode of psychiatric illness, alcohol or drug abuse, major somatic illness or habitual use of nicotine as determined by a health screening carried out by a senior physician. The study and study design were approved by the Regional Ethics Committee in Stockholm and the Karolinska University Hospital Radiation Safety Committee. All subjects gave written informed consent prior to participating.

In addition to the PET examinations, subjects also completed the Swedish universities Scales of Personality (SSP) (12). SSP is an established personality inventory that includes scales measuring social desirability (SocDes) and physical trait aggression (PhTA).

### MRI and PET examinations

Magnetic Resonance Imaging (MRI) and PET examination protocols were similar to those described in our previous study (9). T1-weighted MRI images were acquired for all subjects using a 1.5T Siemens Magnetom Avanto system (Erlangen, Germany). All subjects were examined on a Siemens ECAT HR 47 (CTI/Siemens, Knoxville, TN), with [^11^C]SCH-23390 injected as a rapid bolus (mean injected activity = 327 *±* 40 MBq; mean specific activity = 0.33 *±* 0.19 GBq/*µ*mol; mean injected mass = 388 *±* 237 *µ*g) in the antecubital vein. The whole of striatum (STR) and the sub-region limbic striatum (LST) were selected as regions of interest (ROIs), since they showed strong significant correlations to both SocDes (positive) and PhTA (negative) in our previous study (9). All ROIs were grey-matter masked and automatically delineated on the T1 images using the FMRIB FSL software (13) and the Oxford-GSK-Imanova maximum probability 25% DTI-based atlas. D1-R BP_ND_ values were derived using the simplified reference tissue model with cerebellum as reference region.

### Statistical analysis

Pearson correlation coefficients were calculated between the SocDes (one-sided test expecting a positive direction), PhTA (one-sided test expecting a negative direction) and ROI BP_ND_ values, using an alpha level of 0.05. Since a non-significant p-value in itself does not necessarily mean that a replication attempted failed, a statistical procedure known as *replication Bayes Factor (BF)* (14) was also employed. A replication BF quantifies the strength of evidence in favor of a successful replication (H1), over the null-hypothesis of no correlation (i.e. a failed replication: H_0_). This is done by using the previously published correlation as the prior for H1, and then calculating the predictive adequacy of H_1_ over H_0_. A BF above 3 for H_1_ (BF_10_>3) is commonly interpreted as providing moderate evidence for a successful replication, and a BF above 3 for H_0_ (BF_01_>3) as moderate evidence for a failed replication. A BF above 10 signifies strong evidence in favor of one hypothesis (H_1_ or H_0_), over the other. All statistical modelling was carried out using R (v.3.3.2).

## Results

All subjects’ personality scale scores fell within *±* 2SD of the population for both SocDes and PhTA (see T-scores on the y-axes of Figure 1). Neither of the scales showed a significant relationship to BP_ND_ in the STR or the LST (Figure 1 and Table 1). In fact, replication BF shows that there was strong to moderate evidence for no association between BP_ND_ and SocDes (BF_01_=12.4 for STR, BF_01_=7.2 for LST), compared to the original correlations, and hence signified a failed replication (Figure 2). For PhTA, there was moderate evidence in favor of a failed replication for the LST (BF_01_=3.3, see Figure 2), while the evidence in favor of the null was inconclusive for STR (BF_01_=1.9, see Figure 2).

**Figure 1:**
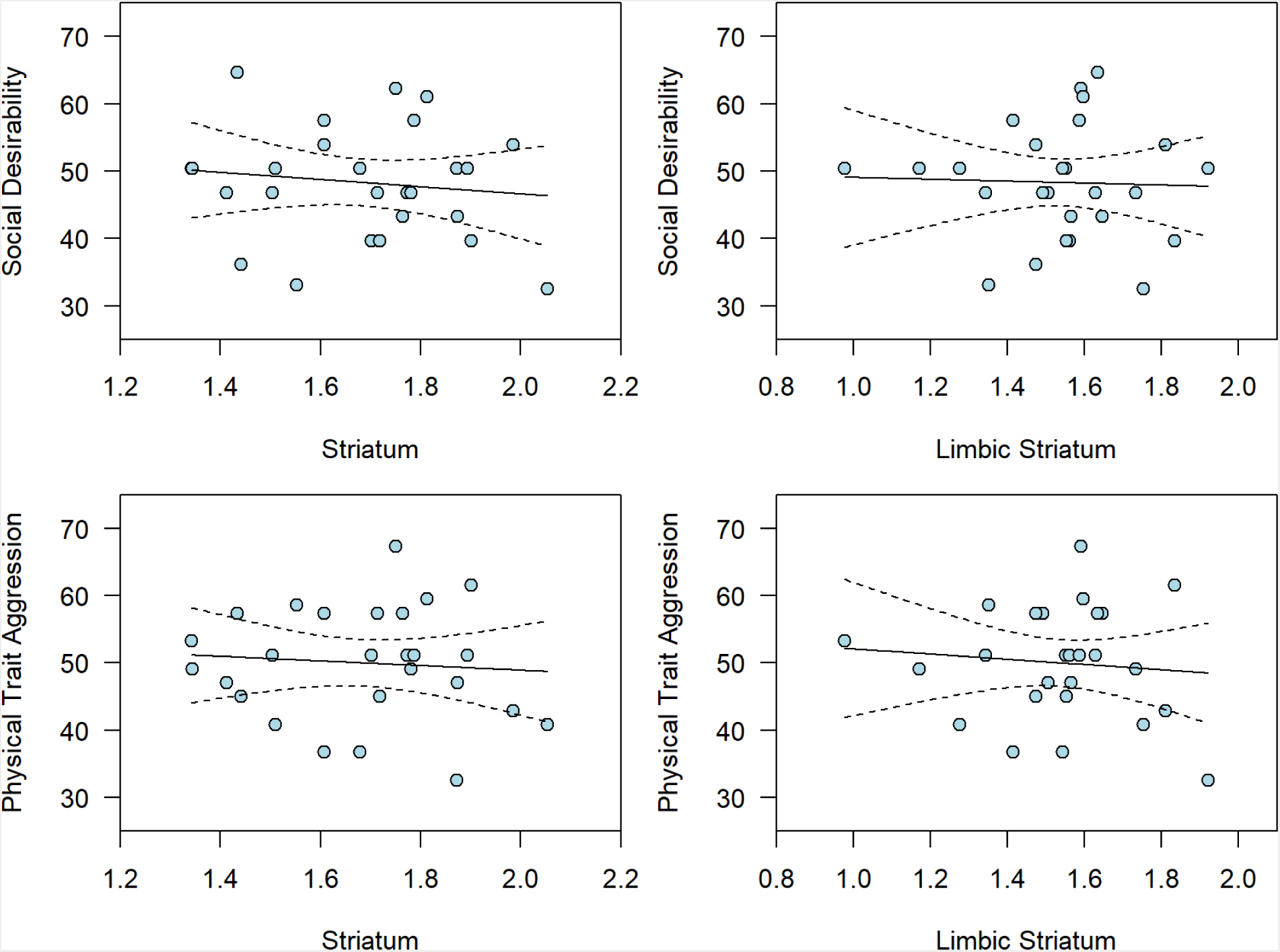
Relationships between D1-R BP_ND_ in striatum and social desirability and physical trait aggression. The dotted lines indicate the 95% confidence intervals. Raw scale scores have been transformed to T-scores (9) for illustrative purposes in this figure.

**Table 1:** Correlations between SocDes and PhTA scores and ROI BPND from the previous (9) and present study. The table also displays the replication BFs which denotes how much support there is for a successful replication, by quantifying how much evidence there is in favor of the original correlation compared to no correlation. Note that the correlation between PhTA and STR was not significant in the original study but have still been included here for completeness.

|  | Original study<sup>a</sup> |  |  | Present study<sup>b</sup> |  |  | Replication BF |  |
| --- | --- | --- | --- | --- | --- | --- | --- | --- |
|  | r | df | p-value | r | df | p-value | BF01 | BF10 |
| **SocDes** |  |  |  |  |  |  |  |  |
| STR | 0.54 | 19 | 0.012 | -0.12 | 24 | 0.73 | 12.4 | 0.08 |
| LST | 0.52 | 19 | 0.015 | -0.03 | 24 | 0.57 | 7.2 | 0.14 |
| **PhTA** |  |  |  |  |  |  |  |  |
| STR | -0.36 | 19 | 0.106 | -0.08 | 24 | 0.35 | 2.0 | 0.51 |
| LST | -0.51 | 19 | 0.019 | -0.09 | 24 | 0.32 | 3.3 | 0.31 |

<sup>a</sup> two-sided test
<sup>b</sup> one-sided test in direction of original study

a two-sided test

b one-sided test in direction of original study

**Figure 2:**
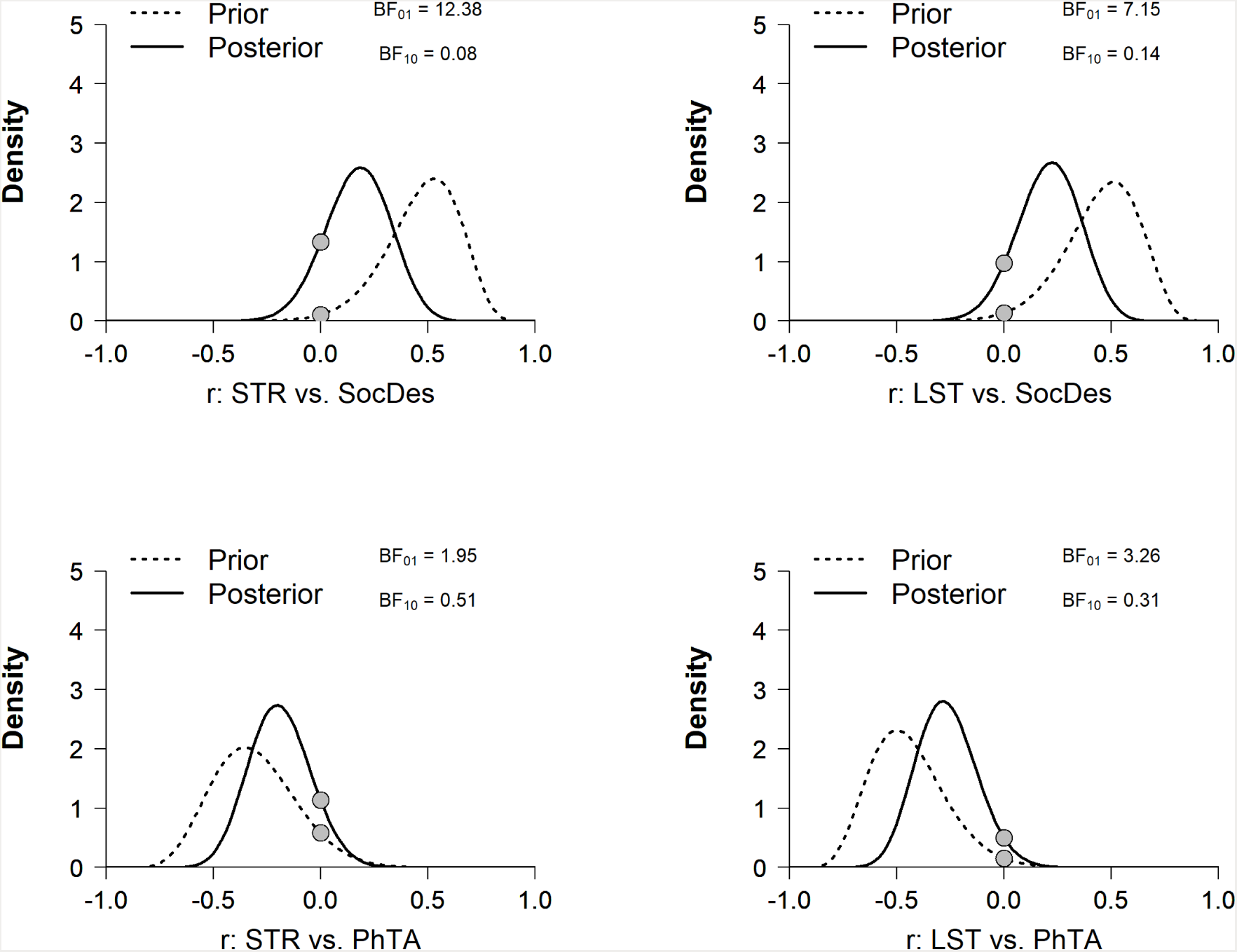
Prior and posterior distributions underlying the replication Bayes factors. In each graph the dotted line denotes the prior which is determined by the correlation from the original study. The posterior (solid line) is obtained by updating the prior using the correlation from the present study. The Savage-Dickey Ratio (the ratio between the heights of the two dots) is then used to calculate the Bayes factor in favour of the original correlation over the null-hypothesis of no correlation. See Verhagen & Wagenmakers (14) for a full explaination of this procedure. In this study, data support the null hypothesis over the original correlations and the Bayes factors hence signifies failed replications.

## Discussion

Using a new and slightly larger sample of healthy subjects, we were not able to replicate our previous findings of an association between D1-R availability in the striatum and social desirability or physical aggression (9). Rather, data showed strong to moderate evidence in favor of failed replications of correlations between D1-R and SocDes or PhTA.

There are several possible explanations for this lack of replication. The present study was based on a sample of healthy young males, while the original study included both males and females from a wider age range. Although both gender and age were controlled for in the original study, the homogeneous sample used in the present study restricts the variance of both D1-R BP_ND_ and the social desirability measures, possibly leading to lower sensitivity to detect an association. Another explanation is that the original findings were false positives, and that there is no direct correlation between D1-R in striatum and pro- and antisocial behavior as measured with SSP. Replication failures are common in science (15, 16). In neuroimaging specifically, small sample sizes and multiple comparisons without adequate correction can lead to incorrect inference. It is also worth noting that a p-value of 0.05, a commonly set threshold for significance, provides only modest evidence in favor of the research hypothesis being true, compared to the null-hypothesis (17).

One way forward is to use a larger and demographically more diverse sample of subjects, in order to maximize both the power and the interindividual variability of PET and personality outcomes. To facilitate this approach, we provide the BP_ND_ and personality data from this study on an online public repository (<u>https://osf.io/te5q7/</u>), so that other PET researchers can pool our data with their samples. Another future line of research could be to correlate D1-R availability with different tests of specific types of pro- and antisocial behavior that could yield more precise outcomes than self-report questionnaires, with larger interindividual variation. Examples of such endophenotypes of social behavior include experimental measures of trustful, altruistic and vindictive decision-making commonly used within the field of behavioral economics (18).

## Author contributions

SC and PPS conceived of the study. PPS designed the study. GJM carried out the image analysis. PPS carried out the statistical analysis. PPS and SC drafted the article. All authors interpreted the results, critically revised the article and approved of the final version for publication.

## Acknowledgement

We thank the staff of the PET group at Karolinska Institutet for their assistance during this study. Code and data can be found at <u>https://osf.io/te5q7/</u> or <u>https://github.com/pontusps/D1R-personality-replication</u>.

